# A second X chromosome improves cognition in aging male and female mice

**DOI:** 10.1101/2024.07.26.605328

**Authors:** Francesca Marino, Dan Wang, Gennifer E. Merrihew, Michael J. MacCoss, Dena B. Dubal

## Abstract

Women show resilience to cognitive aging, in the absence of dementia, in many populations. To dissect sex differences, we utilized the FCG and XY^*^mouse models. Female gonads and sex chromosomes improved cognition in aging mice of both sexes. Further, presence of a second X in male and female mice conferred cognitive resilience while its absence in females blocked it. In the hippocampal proteome of aging female mice, the second X increased proteins involved in synaptogenesis signaling – a potential pathway to improved cognition.

## Main text

The investigation of sex differences and their mechanistic etiologies is of major importance to human health. Understanding what makes one sex more vulnerable or resilient to measures of aging can reveal new targets that may ultimately benefit both men and women.

True sex differences exist in aging. Women live longer than men around the world^1, 2^; they also show resilience to cognitive decline^3-6^ and higher baseline function in typical aging^6-8^ in many populations, when dementia and its subsequent development are carefully excluded^3, 4, 6^. Since cognition is a key manifestation of brain function eroded by aging, understanding sex differences and their causes are high value areas of investigation.

Whether female mice show better cognition in typical aging – and whether sex chromosomes or gonads influence sex difference in cognitive aging – remain largely unknown. To examine this, we utilized genetic mouse models of sex biology^9, 10^. The Four Core Genotype (FCG) model tests whether gonads or sex chromosomes contribute to a sex difference^11^. The XY* model tests whether the X or Y chromosome contributes^12^.

We tested for cognitive aging and any accompanying sex differences in young and aged mice using the two-trial Y maze paradigm (Fig. 1a) which measures spatial and working memory, a target of aging^13-18^. As expected, aging decreased cognition in both sexes (Fig. 1b). While no sex differences were observed in young mice, female sex attenuated age-induced cognitive decline (Fig. 1b).

**Figure 1.**
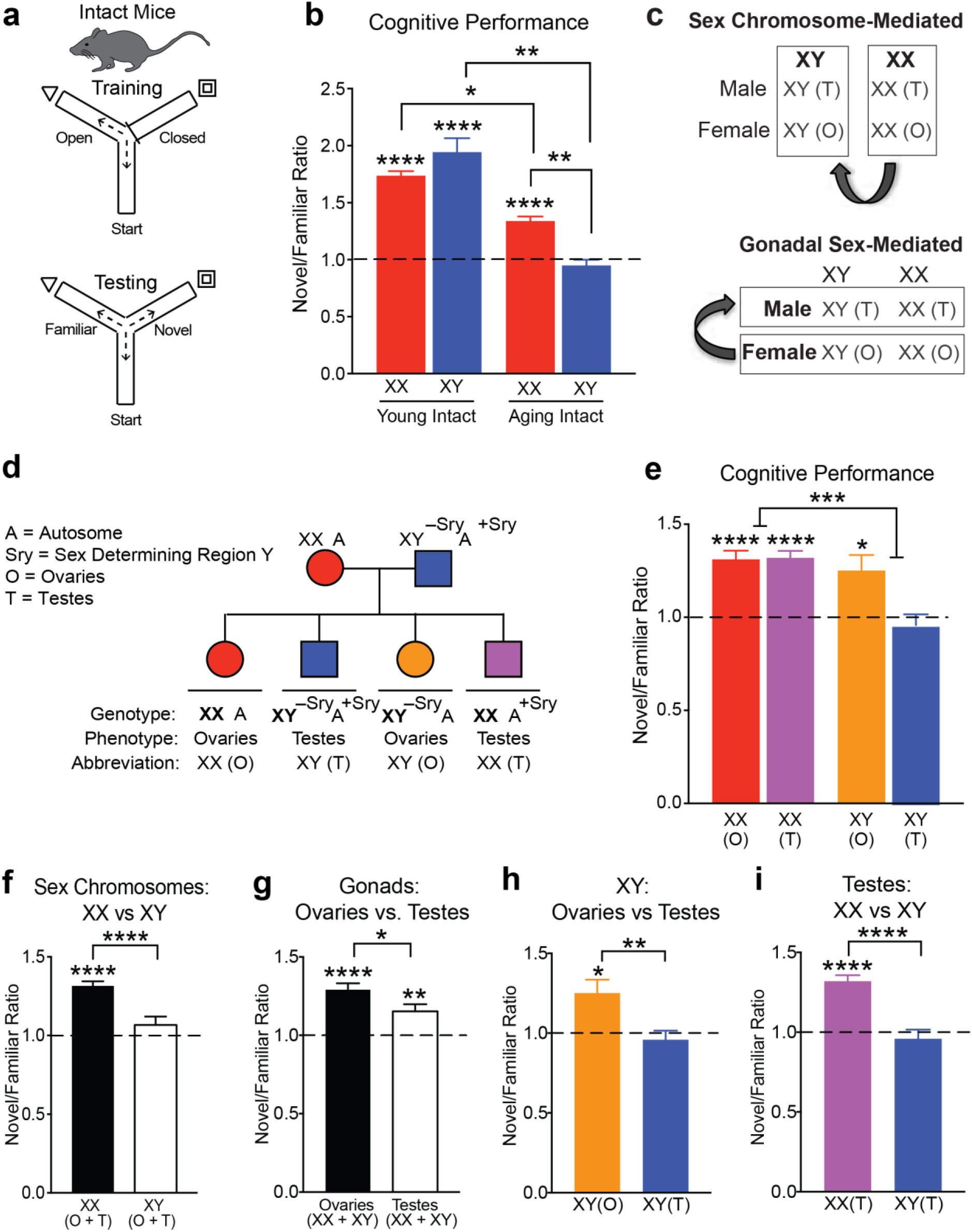
Female sex confers resilience to cognitive aging through sex chromosomal and gonadal effects. **(a)** Schematic of cognitive training and testing in the two-trial Y maze probing spatial and working memory in intact young and aging, male and female mice. **(b)** Ratio of distance traveled in novel arm over familiar arm comparing intact males (XY) with intact females (XX) in the young and old life stages (ages: young 5-7 months, old 19-24 months) (XX young, n = 19; XY young, n = 11; XX old, n = 26; XY old, n = 20). Dashed line represents no learning and memory with a novel/familiar ratio of 1. Aging decreased cognition in both sexes (Two-way ANOVA: aging ****p<0.0001, sex n.s., interaction ****p<0.0001; young XX vs aging XX: *p=0.0160, young XY vs aging XY: **p=0.0320, aging XX vs aging XY: **p=0.0228, Bonferroni corrected two-tailed t-tests). All groups except old XY males showed spatial and working memory (****p<0.0001, two-tailed one sample t-tests vs 1). XX females showed less age-induced cognitive decline compared to XY males (**p=0.0228; Bonferroni). **(c)** Strategy to identify whether sex chromosomes or gonads contribute to sex difference in cognitive aging using the Four Core Genotype (FCG) mouse model. **(d)** Diagram of FCG mouse model. XX females were crossed with XY males with the Sry gene on an autosome instead of the Y chromosome to produce 4 sex genotypes. **(e)** Ratio of distance traveled in novel arm over the familiar arm comparing all four FCG genotypes in an aging FCG cohort (n=68) (age: old 19-22 months) (XX(O), n = 19; XX(T), n = 21; XY(O), n = 10; XY(T), n = 18). Dashed line represents no learning and memory with a novel/familiar ratio of 1. All genotypes except for old typical males (XY, T) showed spatial and working memory. (XX(O) and XX(T): ****p<0.0001, XY(O): *p=0.0150, two tailed one sample t-tests vs 1). Sex chromosomes influenced cognition in aging as the strongest statistical effect (Two-way ANOVA: sex chromosomes ***p=0.0003, gonads *p=0.0110, **interaction p=0.0077). **(f)** XX mice (ovaries + testes) showed better cognition than XY mice (ovaries + testes) in aging. (XX: ****p<0.0001 two-tailed one sample t-test vs 1; ****p<0.0001 two-tailed t-test indicated by brackets). **(g)** Mice with ovaries (XX + XY) showed better cognition than mice with testes (XX + XY) in aging. (Ovaries: ****p<0.0001, Testes: **p=0.0015 two-tailed one sample t-tests vs 1; *p=0.0309 two-tailed t-test indicated by brackets). **(h)** In XY mice, ovaries increased cognition (XY(O): *p=0.0150 two-tailed one sample t-test vs 1; **p=0.0066 two-tailed t-test indicated by brackets). **(i)** In mice with testes, XX sex chromosomes increased cognition (XX(T): ****p<0.0001 two-tailed one sample t-test vs 1; ****p<0.0001 two-tailed t-test indicated by brackets)

Using the FCG mouse model^19^ (Fig. 1c), we began dissecting etiology of the female advantage in cognitive aging. FCG mice harbor a genetic manipulation of the testes determining factor (SRY) that enables generation of XX and XY mice with either ovaries (O) or testes (T): XX(O), XX(T), XY(O), XY(T) (Fig. 1d). In FCG mice, the combination of male sex chromosomes and gonads, XY(T), decreased cognition in aging (Fig 1e), compared to all other groups. XX mice with either ovaries or testes (O+T) showed better cognition compared to XY mice of either gonadal phenotype (O+T), indicating a main effect of sex chromosomes (Fig 1f). Likewise, mice with ovaries (XX+XY) showed better cognition than those with testes (XX+XY), indicating a main effect of gonads (Fig. 1g). Ovaries increased cognition in XY mice (Fig. 1h), and XX chromosomes increased cognition in mice with testes (Fig. 1i). It is important to note that the effect of sex chromosomes persisted despite an insertion of 9 X genes on the Y chromosome in these mice^20^. Collectively, these data show that female genotype (XX) or phenotype (ovaries) rescued cognitive deficits in typical aging male (XY, T) mice.

To further dissect the sex chromosome effect on cognitive aging, we examined sex differences in cognition of young and aging mice – in the absence of gonads. To this goal, we gonadectomized (Gnx) young and aging mice following sexual differentiation and maturity and assessed their cognition using the two-trial Y maze (Fig. 2a). While no sex differences were observed in Gnx young mice, female sex (XX) continued to attenuate age-induced cognitive decline (Fig. 2b). Thus, “typical” aging females (XX) continued to show resilience to cognitive decline compared to “typical” aging males – even without gonadal hormones. These data further suggest a role for sex chromosomes in the sex difference.

**Figure 2.**
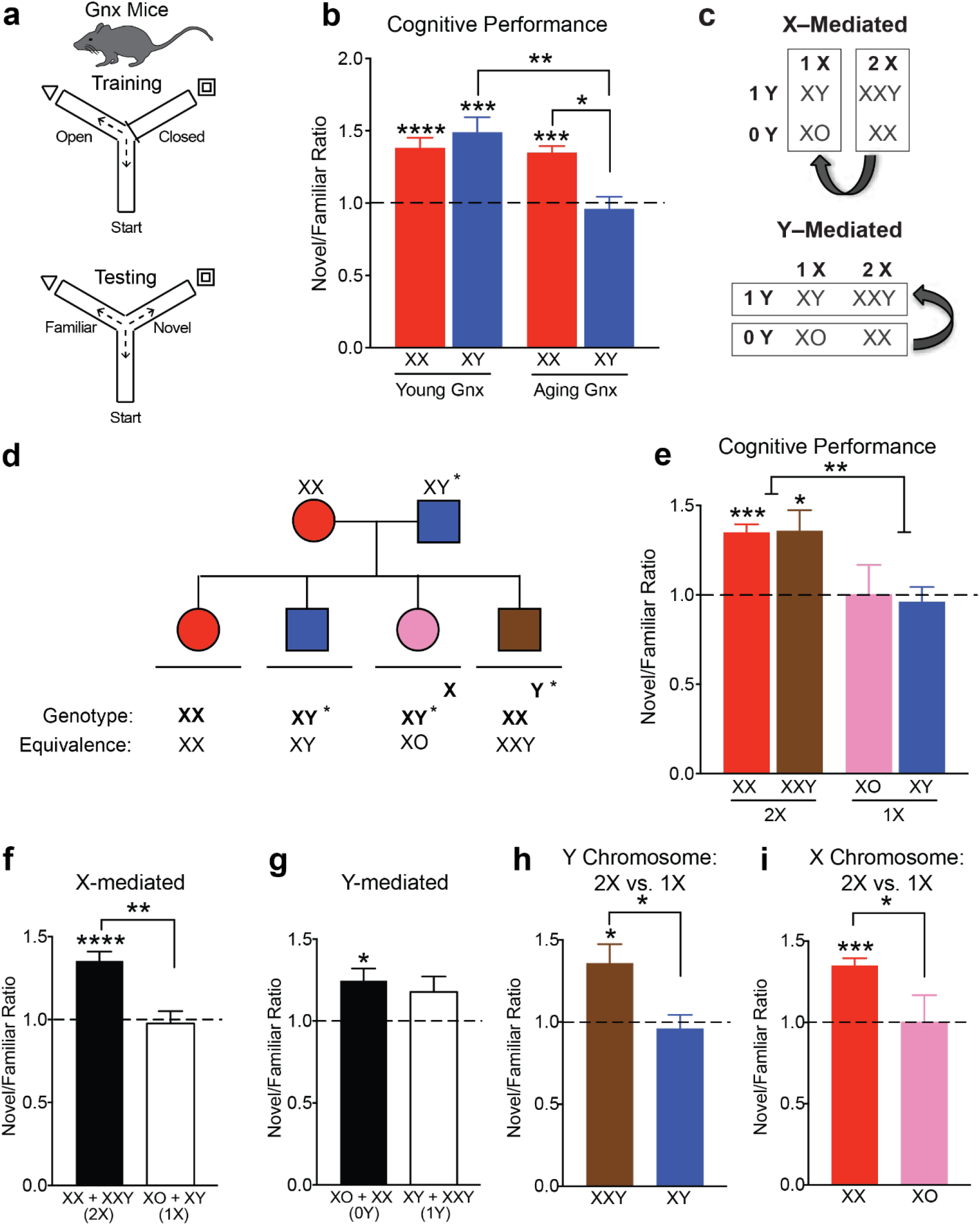
A second X chromosome confers resilience to cognitive aging in male and female mice. **(a)** Schematic of cognitive training and testing in the two-trial Y maze probing spatial and working memory in gonadectomized (Gnx) young and aging, male and female mice. **(b)** Ratio of distance traveled in novel arm over familiar arm comparing Gnx males (XY) and Gnx females (XX) in the young and old life stages (ages: young 5-7 months, old 22-24 months) (XX young, n = 14; XY young, n = 8; XX aged, n = 7; XY young n = 5). Dashed line represents no learning and memory with a novel/familiar ratio of 1. Aging decreased cognition in old, Gnx males (Two-way ANOVA: aging **p=0.0032, sex n.s., interaction **p=0.0080; **p=0.0030, Bonferroni corrected two-tailed t-test). All groups except old, Gnx XY males showed spatial and working memory (young XX: ****p<0.0001, young XY: ***p=0.0021, aging XX: ***p=0.0002, two tailed one sample t-tests vs 1). Gnx, XX females showed less age-induced cognitive decline compared with Gnx, XY males (*p=0.05, Bonferroni). **(c)** Strategy to identify whether the sex chromosome effect depends on the X or Y chromosome using the XY* mouse model. **(d)** Diagram of XY* mouse model. XX females were crossed with XY* males that harbor an altered pseudoautosomal region on the Y chromosome, allowing abnormal crossover with the X chromosome during meiosis. **(e)** Ratio of distance traveled in novel arm over the familiar arm comparing all four XY* genotypes in a Gnx, aging XY* cohort (n=21) (age: old 22-24 months) (XX, n=7; XXY, n=6: XO, n=3; XY, n=5) Dashed line represents no learning and memory with a novel/familiar ratio of 1. All genotypes except for typical males (XY) and XO females showed spatial and working memory (XX: ***p=0.0002, XXY: *p=0.0268, two tailed one sample t-tests vs 1). Increased dose of X chromosomes improved cognition (Two-way ANOVA: X chromosome **p=0.0015, Y chromosome n.s., interaction n.s.). **(f)** Mice with two X chromosomes (XX+XXY) showed better cognition than mice with one X chromosome (XY+XO) (****p<0.0001, two-tailed one sample t-test vs 1) (**p=0.0006, two-tailed t-test indicted by bracket). **(g)** Mice with no Y chromosome (XX+XO) showed similar cognition compared to mice with one Y chromosome (XY+XXY) (*p=0.0090, two-tailed one sample t-test vs 1). **(h)** A second X chromosome increased cognition in male mice (XXY vs XY) (XXY: *p=0.0268, two-tailed one sample t-test vs 1; *p=0.0125, two-tailed t-test indicated by bracket), **(i)** A second X chromosome increased cognition in female mice (XX vs XO) (XX: ***p=0.0002, two-tailed one sample t-test vs 1; *p=0.0206, two-tailed t-test indicated by bracket)

Next, we dissected the effects of the X and Y chromosome on cognitive aging using the XY* model^21^. XY* males harbor an altered pseudoautosomal region on the Y chromosome, allowing for abnormal crossover with the X during meiosis which leads to the generation of XX, XY, XXY, and XO mice (Fig. 2d). A sex difference that varies by the presence of a Y is Y-mediated and one that varies by the dosage of the X is X-mediated (Fig. 2c). In Gnx, aging XY* mice, XO and XY mice showed deficits in cognition (Fig. 2e). Indeed, mice with two X chromosomes (XXY+XX) showed better cognition than those with one X chromosome (XY+XO), indicating a main effect of X chromosome dosage (Fig. 2f). In contrast, mice with a Y chromosome (XY+XXY) or without it (XX+XO) showed similar cognition, indicating that the Y did not decrease cognition in aging (Fig. 2g). Specifically, a second X chromosome increased cognition in male (Fig. 2h) and female (Fig. 2i) mice. Therefore, a second X chromosome conferred resilience to cognitive aging.

Since a second X chromosome improved cognition in aging, we specifically probed whether its presence – compared to its absence – alters the hippocampal, aging proteome. To more precisely isolate the effect of the second X in the absence of the Y chromosome and gonadal hormones, we executed the study in Gnx, littermate XX compared to XO female mice.

A proteomic analysis of aged XX and XO mice identified 3692 unique proteins conserved across genotypes. From this total, 124 X chromosome proteins were detected out of the 1747 known X chromosome proteins in Mus musculus^22, 23^. Across the genome, 103 differentially expressed proteins were detected (Fig. 3a). Seven X chromosome proteins were differentially expressed between XX and XO mice: Atp6ap2, Dlg3, Nlgn3, Apool, Rbm41, Frmpd4, and Hccs (Fig. 3b). The latter two were upregulated in young and aging, intact and Gnx XO mice (Fig 3a, Extended Fig. a-d) in the XY* line likely due to an insertion^11, 24^. To our knowledge, this is the first report that the second X increased or decreased the other five X chromosome-derived proteins – collectively involved in synaptic plasticity, RNA Binding, ATP, and mitochondria – in the aging hippocampus.

**Figure 3.**
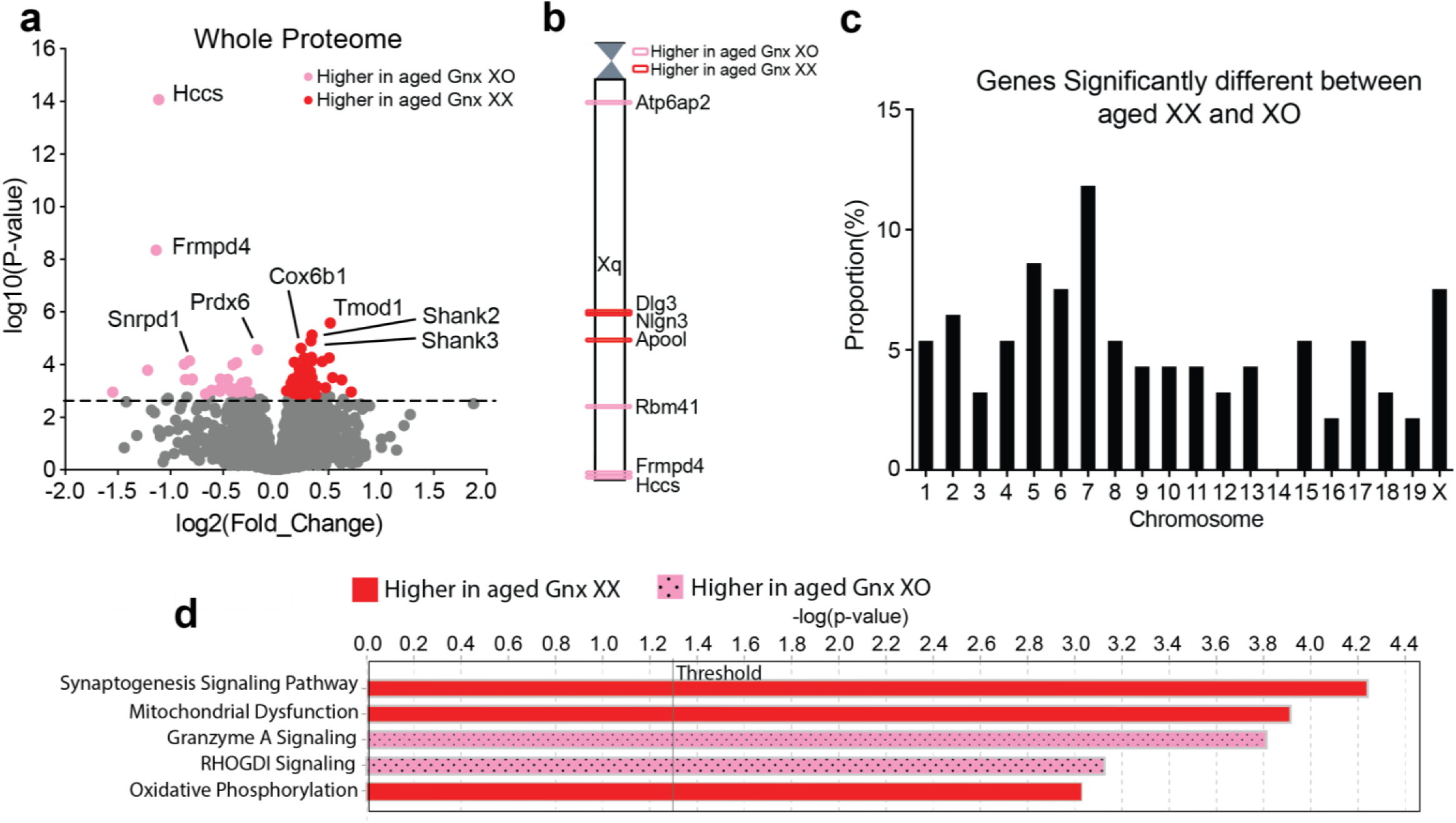
A second X chromosome influences protein expression across the genome. **(a)** Proteomics analysis of hippocampi from aged Gnx female mice (n=4 XO; n=6 XX) (age: 22-24 months). Volcano plot of protein expression comparing aged Gnx XO and aged Gnx XX females. Colored circles indicate significant differentially expressed proteins with a –log10(p-value) threshold of 2.8 which corresponds to an FDR value of 0.052. 103 significant proteins were detected, and the top eight differentially expressed proteins are labeled. **(b)** Chromosome map displaying the location of X chromosome genes corresponding to significant differentially expressed X proteins. Color indicates the genotype with higher protein expression. **(c)** Bar plot of the proportion of genes corresponding to significant differentially expressed proteins by chromosome number. A second X chromosome influences protein expression across the genome. **(d)** Ingenuity pathway analysis highlighting the top biological pathways that are significantly different between aging XX and XO females. A second X increases synaptogenesis signaling. Threshold indicates a –log10(p-value) of 1.3 which corresponds to a p-value of 0.05. Color indicates the genotype with higher pathway expression.

Interestingly, the presence of a second X chromosome caused proteins to be differentially expressed in aging from genes represented on nearly every chromosome (Fig. 3c). More proteins were upregulated in aging XX mice (76 proteins) compared to those in XO mice (27 proteins). Furthermore, a functional analysis of these differentially expressed proteins with IPA revealed that a second X chromosome increased proteins involved in the synaptogenesis signaling pathway (Fig. 3d), suggesting that synapses may be central to mechanisms of cognitive resilience in XX mice. In an additional experiment of young Gnx XX and XO mice, IPA analysis revealed that a second X increases the myelination signaling pathway (Extended Data Fig. 1e), suggesting that boosting myelination early on may be important to decreasing cognitive deficits later in life.

Our studies collectively revealed that mice with two X chromosomes show resilience to cognitive decline in aging regardless of gonadal sex or the presence of a Y chromosome.

Furthermore, a second X chromosome influenced protein expression across the proteome and increased synaptogenesis signaling and myelination pathways. These data support the hypothesis that XX females are more resilient to cognitive decline in typical aging.

The second X chromosome altered proteins coding from genes across nearly all autosomes in our mouse studies. Notably, individuals with Turner’s syndrome, who partially or fully lack a second X, show global hypomethylation of their genome, and similarly exhibit altered gene expression of autosomes^25^. In addition, genes on their single X chromosome can be differentially methylated^26^ and expressed, suggesting human relevance for our proteomics analysis revealing differences between XX and XO mice on both autosomes and the X chromosome. It is interesting to speculate that the presence of a second X chromosome could increase broadly acting methylation factors upstream of genomic transcription. Alternatively, the absence of a Barr body, the highly condensed chromatin structure surrounding the inactive X, could physically change the transcriptional landscape of autosomes and the active X chromosome.

Potential limitations of our study include that mice were tested in one cognitive task and FCG mice harbored an insertion of 9 X genes on the Y chromosome^20^. We used the two-trial Y maze, including training and testing, because it probes both spatial and working memory^27^ and has been particularly sensitive to measuring cognitive deficits in our studies of aged mice and disease models^13, 14, 16^. Since it utilizes primarily exploration, stress likely contributed less to cognitive measures compared to other paradigms. Despite the insertion of 9 X genes in FCG XY males^20^, which could dilute a sex chromosomal effect, XX mice continued to show better cognition that XY mice, regardless of gonads. In support of these findings, the X chromosome effect was further probed and demonstrated in XY* mice, an independent transgenic mouse model.

Strategies to target and recapitulate XX-mediated cognitive resilience in the brain could lead to development of novel therapies to treat deficits in aging males, females, or both.

## Methods

### Animals

Mice for in vivo and proteomics studies were on a congenic C57BL/6J background and kept on a 12-hour light/dark cycle with *ad libitum* access to food and water. The standard housing was 5 mice per cage based on gonadal sex. Two-trial Y maze studies were carried out during the light cycle. All animal studies were approved by the Institutional Animal Care and Use Committee of the University of California, San Francisco and conducted in compliance with NIH guidelines. All mice included in this study were bred from the Fore Core Genotypes model^19^ and XY* model^21^. Intact and gonadectomized mice were used as indicated. For hormonal depletion, male and female mice underwent gonadectomy following sexual differentiation and after reaching reproductive maturity around 75 days. Behavioral and proteomics experiments were conducted in a blinded manner during data collection and analysis, unless otherwise indicated.

### Two-Trial Y Maze

Mice were tested on the two-trial Y maze (Noldus Ethovision) to assess spatial and working memory as described^15-18^. Briefly, mice were trained by exploring the maze with a visual cue in one arm and another arm blocked off. 16 hours after this training session, mice underwent testing with all three arms open (start arm, familiar arm, novel arm) and the distance travelled exploring the novel arm compared to the familiar arm was measured. This ratio of novel to familiar arm distance represents an index of memory.

### Proteomics

A total of 46 mouse hippocampus samples were used for proteomics. Mice were from two genotypes (XX and XO), two age groups (young: 4-6 months and old: 24-26 months), and two ovary conditions (intact and ovariectomized (GNX)). All mice were female. Samples were randomized and processed in 4 batches. In addition, one hippocampus specific reference and one brain reference were included in each batch as quality controls.

Mouse hippocampus tissue was originally resuspended in 90 μl of 0.1% RapiGest, 50 mM ammonium bicarbonate and 1X HALT phosphatase and protease inhibitors, vortexed and briefly sonicated at setting 3 for 10 s with a Fisher sonic dismembrator model 100. Sample buffer was then adjusted to 5% SDS, 50 mM triethylammonium bicarbonate (TEAB), 2 mM MgCl2 and 1X HALT phosphatase and protease inhibitors for S-trap column digestion (Protifi)^28^. Demultiplexed DIA data was analyzed using EncyclopeDIA (version 0.6.14) to align the quantitative DIA data to the gas-phase fractionated DIA data using the Uniprot mouse canonical FASTA (March 2018). The quantitative spectral library was imported into Skyline-daily 4.2.1.19058 with the UniProt mouse canonical FASTA as the background proteome to map peptides to proteins^28^. The mzML data is imported and all data is TIC normalized.

The TIC-normalized data was preprocessed using the MSstats package^29^, eliminating peptides that appeared in multiple proteins and proteins containing only a single feature. In this context, a feature refers to a combination of PeptideSequence, PrecursorCharge, FragmentIon, and ProductCharge. The resulting dataset contained 3694 proteins common between all four batches. To impute left-censored missing data, we employed a Quantile Regression Imputation of Left-Censored Data (QRILC) approach using the R package imputeLCMD^30, 31^. Given that our biological unit of interest was protein, we aggregated features into protein-level summaries. This step was executed using the Tukey’s Median Polish (TMP) summary method within MSstats’ dataProcess function. To remove the unwanted data variations, the Bioconductor SVA package was utilized to identify and adjust for three surrogate variables (SVs) as covariates in later analyses^32^. To identify group differences at the protein level, a linear model was fitted using the Bioconductor limma package^33^, incorporating the estimated SVs as covariates in the model. The limma package uses empirical Bayes moderated statistics, which improves power by ‘borrowing strength’ between proteins in order to moderate the residual variance^34^. We applied the Benjamini-Hochberg procedure for multiple testing correction. Proteins were sorted for significant differential abundance using a cutoff of less than 10% FDR. Bar graphs and volcano plots were created in R with the ggplot2 and ggrepel packages. Functional biological pathway analysis was performed with Qiagen IPA (Summer Release 2023).

### Statistics

GraphPad Prism (version 7.0) was used for *t* tests and visualization of data. R (nmle package) was used for analysis of variance (ANOVAs) and Bonferroni post hoc tests. Differences between two means were assessed by 2-tailed *t* tests unless otherwise indicated. Differences between multiple means were assessed by 2-way ANOVA. Multiple comparisons of post hoc *t* tests were corrected with the Bonferroni-Holm test. Exclusion criteria (greater than 2 SDs above or below the mean) were defined a priori to ensure unbiased execution of outliers. Error bars represent mean ±SEM. Null hypotheses were rejected at or below a *p* value of 0.05.

## Data availability

All behavior data supporting the findings of this paper are available within the source data file provided with this paper. The Skyline documents and raw files for the proteomics DIA MS/MS data are available at Panorama Public^35^ ProteomeXchange ID: PXD049347. Access URL: https://panoramaweb.org/mouse-X-chromosome-cognitive-aging.url.

## Code availability

No new code was created for the data in this manuscript.

## Acknowledgements

We thank Chen Chen, Emily Davis, and Arturo Moreno for technical assistance with mouse behavioral studies and Arthur Arnold for discussions. Supported by NIH grants AG034531, AG038325 (D.B.D.), AG065156, AG013280 (MM), the Simons Foundation (D.B.D.), Bakar Aging Research Institute (BARI) (FM and D.B.D) and philanthropy (D.B.D). Funders had no role in study design, data collection, analysis, decision to publish or preparation of the manuscript.

## Author Contributions

F.M., G.E.M, D.W., M.M., D.B.D. carried out experimental design and studies, analyzed data and/or contributed to the manuscript. All authors discussed the results and commented on the manuscript.

## Competing Interest Statement

F.M., G.E.M., D.W., M.M., D.B.D. have no competing interests related to the study.

## Figure Legend (for Extended Figure)

**Extended Data Figure 1.**
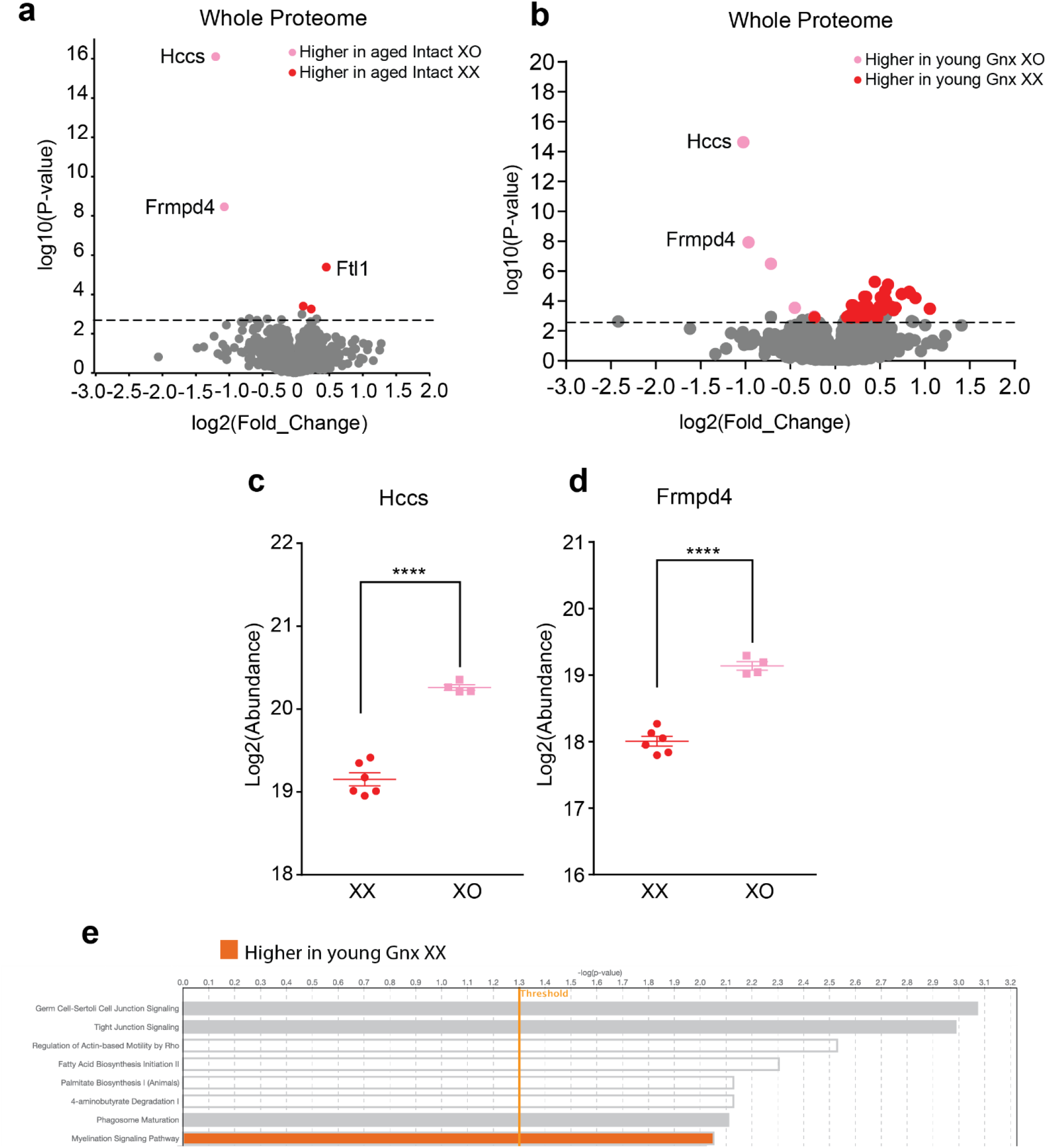
**(a)** Proteomics analysis of hippocampi from aged intact female mice (n=6 XO; n=6 XX) (age: 22-24 months).Volcano plot of protein expression comparing aged Gnx XO and aged Gnx XX females. Colored circles indicate significant differentially expressed proteins with a –log10(p-value) threshold of 2.8 which corresponds to an FDR value of 0.052. 5 significant proteins were detected and are labeled. **(b)** Proteomics analysis of hippocampi from young Gnx female mice (n=6 XO; n=6 XX) (age: 4-6 months). Volcano plot of protein expression comparing aged Gnx XO and aged Gnx XX females. Colored circles indicate significant differentially expressed proteins with a –log10(p-value) threshold of 2.8 which corresponds to an FDR value of 0.052. 60 significant proteins were detected, and the top eight differentially expressed proteins are labeled. **(c)** Scatter plot of protein expression for the highest differentially expressed protein: Hccs. (****p<0.0001 two-tailed t-test indicated by bracket). **(d)** Scatter plot of protein expression for the second highest differentially expressed protein: Frmpd4. (****p<0.0001 two-tailed t-test indicated by bracket). **(e)** Ingenuity pathway analysis highlighting the top biological pathways that are significantly different between young Gnx XX and XO females. A second X increases myelination signaling pathway. Threshold indicates a –log10(p-value) of 1.3 which corresponds to a p-value of 0.05. Color indicates the genotype with higher pathway expression.

## Notes

https://panoramaweb.org/mouse-X-chromosome-cognitive-aging.url

### Competing Interest Statement

The authors have declared no competing interest.

